# RNA processing/modifying enzymes play key roles in the response to thermospermine in *Arabidopsis thaliana*

**DOI:** 10.1101/2022.09.19.508594

**Authors:** Takahiro Tanaka, Daiki Koyama, Mitsuru Saraumi, Hiroyasu Motose, Taku Takahashi

## Abstract

Thermospermine acts in negative regulation of xylem differentiation through enhancing mRNA translation of the members of the *SAC51* gene family in Arabidopsis. These mRNAs contain conserved upstream open-reading-frames (uORFs) that are inhibitory to the main ORF translation. To address the mode of action of thermospermine in this process, we have isolated mutants that are insensitive to thermo<u>s</u>permine, named *its*. We show here that four genes responsible for the mutants, *its1* to *its4*, encode a homologue of SPOUT RNA methyl transferase, an rRNA pseudouridine synthase CBF5/NAP57, a putative spliceosome disassembly factor STIPL1/NTR1, and a plant-specific RNA-binding protein PHIP1, respectively. While these mutants except *its1* are almost normal in appearance, they enhance the dwarf phenotype of a mutant of *ACL5* defective in thermospermine biosynthesis, resulting in tiny-sized plants reminiscent of a double knockout of *ACL5* and *SACL3*, a member of the *SAC51* family. We confirmed that the GUS reporter activity from the *SAC51* 5’-GUS fusion transcript was severely reduced in all of these mutants. These results unveil the importance of RNA processing and modification for the translation of transcripts containing regulatory uORFs.

## Introduction

Thermospermine is produced from spermidine and an aminopropyl donner, decarboxylated S-adenosyl methionine, by thermospermine synthase (Knott et al., 2007) and ubiquitously detected in the plant kingdom (Minguet et al., 2008; Takano et al., 2012; Solé-Gil et al., 2019). In *Arabidopsis thaliana*, the *ACAULIS5* (*ACL5*) gene encoding thermospermine synthase is specifically expressed in xylem precursor cells and required for negative regulation of xylem differentiation (Clay and Nelson, 2005; Kakehi et al, 2008; Muñiz et al., 2008). Loss-of-function mutants of *ACL5* have excess xylem vessels and show dwarfed growth (Hanzawa et al., 1997). The phenotype is partially recovered by exogenous thermospermine or a structurally related tetraamine, norspermine (Kakehi et al., 2010). In contrast, the growth of wild-type seedlings in the presence of thermospermine results in the repression of xylem differentiation and lateral root formation (Tong et al., 2014). Studies of suppressor mutants that restore the dwarf phenotype of *acl5*, have identified *SUPPRESSOR OF ACL5* (*SAC*) genes and revealed that thermospermine enhances mRNA translation of *SAC51*, which encodes a basic helix-loop-helix (bHLH) protein (Imai et al., 2006). The *SAC51* mRNA contains upstream open-reading frames (uORFs) in the 5’ leader sequence, one of which is highly conserved among different plant species and acts to interfere with the main ORF translation (Hayden and Jorgensen, 2007; Tran et al., 2008; Jorgensen and Dorantes-Acosta, 2012; von Arnim et al., 2014). Mutations in this uORF behave as a dominant suppressor of *acl5* and similar uORF mutations have been also identified in the members of the *SAC51* gene family, *SACL1* and *SACL3* (Vera-Sirera et al., 2015; Cai et al., 2016) We have further revealed that other suppressor mutants of *acl5, sac52-d, sac53-d*, and *sac56-d*, are dominant alleles of *RPL10A, RACK1A*, and *RPL4A*, encoding a ribosomal protein L10, an integral ribosomal component Receptor for Activated C Kinase1, and a ribosomal protein L4, respectively (Imai et al., 2008; Kakehi et al., 2015). These three ribosomal mutations have been suggested to suppress the *acl5* phenotype by attenuating the inhibitory effect of the uORFs on the translation of the main ORF of *SAC51* family genes.

The *ACL5* expression is induced by auxin (Hanzawa et al., 2000). Mechanistically, heterodimers of bHLH transcription factors, LONESOME HIGHWAY (LHW) and TARGET OF MONOPTEROS5 (TMO5) or TMO5LIKE1 (T5L1) activate the *ACL5* expression in xylem precursor cells of the root (Katayama et al., 2015). *TMO5* and *T5L1* are direct targets of an auxin responsive transcription factor ARF5/MONOPTEROS (Donner et al., 2009; Schlereth et al., 2010). LHW-TMO5 and LHW-T5L1 heterodimers also activate expression of *SACL3*, a cytokinin biosynthetic gene *LONELY GUY4* (*LOG4*), and *ARABIDOPSIS HISTIDINE PHOSPHOTRANSFER PROTEIN6* (*AHP6*), which encodes a cytokinin signaling inhibitor (De Rybel et al., 2014; Ohashi-Ito et al., 2014; Katayama et al., 2015; Vera-Sirera et al., 2015). SACL3 in turn acts to form a heterodimer with LHW antagonistically to TMO5 and T5L1, and consequently functions as a repressor of both the *ACL5* expression and cytokinin-dependent provascular cell division (Katayama et al., 2015; Vera-Sirera et al., 2015). Our previous studies using the GUS reporter gene fusions have suggested that the main ORF translation of *SAC51* and *SACL1* is significantly enhanced by thermospermine but that of *SACL3* and *SACL2* is not (Cai et al., 2015). In contrast to an overexpression allele *sac51-d*, a loss-of-function allele of *SAC51, sac51-1*, has no additional effect on the *acl5* phenotype but that of *SACL3, sacl3-1*, severely enhances the *acl5* phenotype, resulting in tiny-sized plants. Moreover, the *sac51-1 sacl3-1* double mutant has an increased level of thermospermine due to an increased expression of *ACL5* and shows an insensitivity to exogenous thermospermine but no morphological phenotype (Cai et al., 2015). These results suggest that *SAC51* and *SACL3* are functionally redundant in the control of *ACL5* expression but *SACL3* has a role independent of the presence of thermospermine. The interaction target(s) of SAC51 and the functional importance of SACL1 and SACL2 remain unknown.

In the present study, aiming at clarifying the precise mode of action of thermospermine from transport to perception and in the regulation of specific mRNA translation, we screened for thermospermine-insensitive mutants of Arabidopsis that can grow at high concentrations of thermospermine. Our results unexpectedly show that all the four genes responsible for these mutants so far identified encode proteins related to RNA processing or modification, providing evidence for critical roles of RNA metabolism in the response to thermospermine.

## Results

### Isolation of thermospermine-insensitive mutants

In the presence of 100 μM thermospermine, wild-type Arabidopsis seedlings show severely reduced growth of the shoot with a reduction of xylem development and few lateral roots. Based on these morphological responses, approximately 10,000 M2 seedlings derived from ethyl methane sulfonate-mutagenized seeds were screened for mutants whose growth is not or less affected by thermospermine. We identified 11 putative mutants, which were later found to include identical alleles by genome mapping and sequencing and integrated into five alleles of four loci named *insensitive to thermospermine1-1* (*its1-1*), *its1-2, its2, its3*, and *its4*.

As shown in Figure 1A, the growth of 10-day-old seedlings of these mutants is less affected by 100 μM thermospermine than that of the wild type. Measurement of the length of the first foliage leaf indicated that *its1, its2*, and *its4* have a similar insensitivity to thermospermine but *its3* is slightly weaker than others (Figure 1B). These mutants also show the insensitivity to thermospermine in terms of the number of lateral roots (Figure 1C). On the other hand, the length of the taproot is unaffected by thermospermine in both the wild type and these mutants (Figure 1D). When grown without thermospermine, *its1* seedlings have hypocotyls with an obviously thicker central cylinder than have the wild-type seedlings, a phenotype mimicking thermospermine-defective *acl5* (Figure 1E). We have examined the effect of thermospermine on ectopic xylem differentiation from mesophyll cells in cotyledons, which is induced by concomitant treatment with auxin, cytokinin, and a strong activator of brassinosteroid (BR) signaling, bikinin. This is according to an in vitro culture system, VISUAL, in which Arabidopsis leaf disks or detached cotyledons cultured with these three plant growth regulators develop xylem vessel elements and phloem sieve cells ectopically in mesophyll cells (Kondo et al., 2016). In our study, intact seedlings were simply grown with these compounds. Wild-type and all *its* mutant seedlings similarly have ectopic xylem vessels around veins especially at the tip of cotyledons (Figure 1F). Cotreatment with thermospermine resulted in narrowed veins and severely repressed ectopic xylem vessel formation in wild-type and *its3* seedlings but had no such effect on *its1, its2*, and *its4* seedlings (Figure 1F).

**Figure 1.**
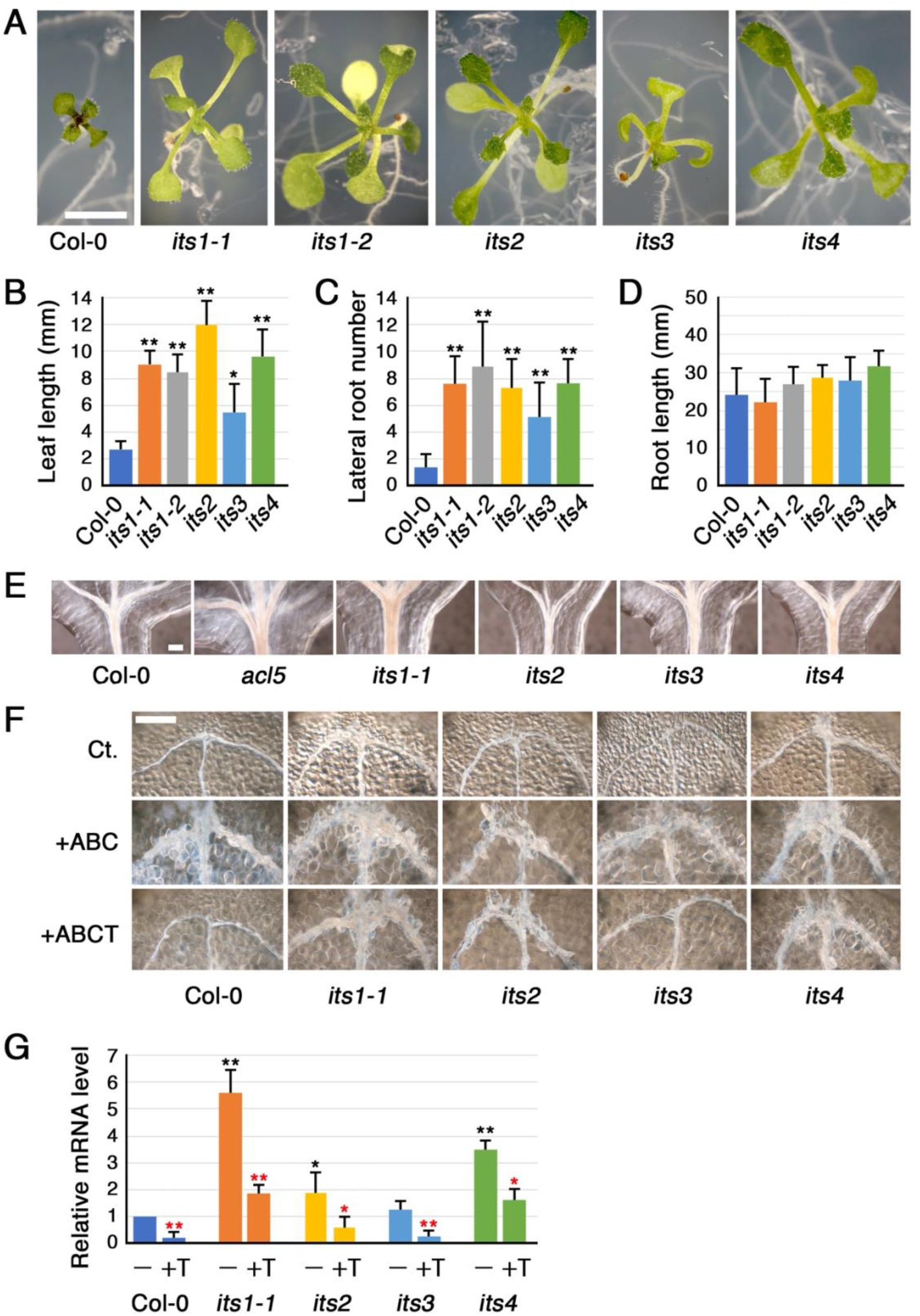
Phenotypes of thermospermine-insensitive mutants. A, Phenotypes of 7-day-old seedlings of the wild type (Col-0), *its1-1, its1-2, its2, its3*, and *its4* grown with 0.1 mM thermospermine. Scale bar, 5 mm. B-D, Comparison of the length of the first leaf (B), the number of lateral roots (C), and the length of the taproot (D) between Col-0 and mutant seedlings grown for 7 days with 0.1 mM thermospermine. Values are means ± SE (n = 10). Asterisks indicate significant difference from Col-0 (\**P* < 0.05; \*\**P* < 0.01, Two-sided Student’s *t* test). E, Vascular phenotypes in hypocotyls of Col-0 and mutant seedlings grown for 7 days without thermospermine. Samples were cleared with chloral hydrate. Scale bar, 200 μm. F, Effect of thermospermine on ectopic xylem vessel differentiation in cotyledons of Col-0 and mutant seedlings. 4-day-old seedlings grown in agar plates were transferred to liquid media without (Ct) or with auxin, bikinin, and cytokinin (+ABC) or also with thermospermine (+ABCT) for further 4 days. Samples were cleared with chloral hydrate. Scale bar, 200 μm. G, Effect of thermospermine on the expression of *ACL5*. Each seedling was incubated for 24 h without (–) or with thermospermine (+T). mRNA levels were normalized to the *ACT8* mRNA level and set to 1 in Col-0 (–). Values are means ± SE (n = 6). Asterisks in black indicate significant difference between Col-0 (–) and each mutant (–) and asterisks in red indicate that between – and +T seedlings (\**P* < 0.05; \*\**P* < 0.01, Two-sided Student’s *t* test).

Quantitative RT-PCR experiments revealed that the steady-state level of *ACL5* expression is significantly higher in *its1, its2*, and *its4* seedlings than in wild-type and *its3* seedlings but it is down-regulated by one-day treatment with thermospermine in all the mutants (Figure 1G), indicating that these mutants retain the ability to respond to thermospermine at the molecular level.

### *ITS1* encodes a homologue of SPOUT1

Genome mapping and sequence analysis revealed that two *its* mutants represent different alleles of a gene, At5g19300, encoding a homologue of the human SPOUT domain-containing methyltransferase 1 (SPOUT1), also designated as C9orf114. C9orf114 was reported to interact with mRNA (Baltz et al., 2012; Castello et al., 2012) while it was also shown to bind to a specific miRNA and its knockdown causes a reduced level of the mature miRNA, suggesting a role in posttranscriptional regulation (Treiber et al., 2017). The yeast orthologs have been suggested to play a key role in the formation of the large ribosomal subunit (Ismail et al., 2022). Because At5g19300 has not been characterized before, we name this locus *ITS1* and refer to one mutant resulting in an amino acid substitution as *its1-1* and another one containing a premature stop codon as *its1-2* (Figure 2A). The Cys-to-Tyr substitution in *its1-1* is present in a region conserved among plants and animals (Figure 2B). We further confirmed that a T-DNA insertion mutant of *ITS1*, which we name *its1-3*, also shows an insensitive phenotype to thermospermine in terms of leaf growth and all these three alleles behave as recessive mutations (Figure 2C). Under standard growth conditions, veins of cotyledons are also thicker in *its1-3* than in the wild type (Figure 2D). Although the first leaf of *its1* seedlings has normal length, their adult flowering plants have a slightly reduced plant height with almost half size of rosette leaves, compared with the wild type (Figure 2, E and F).

**Figure 2.**
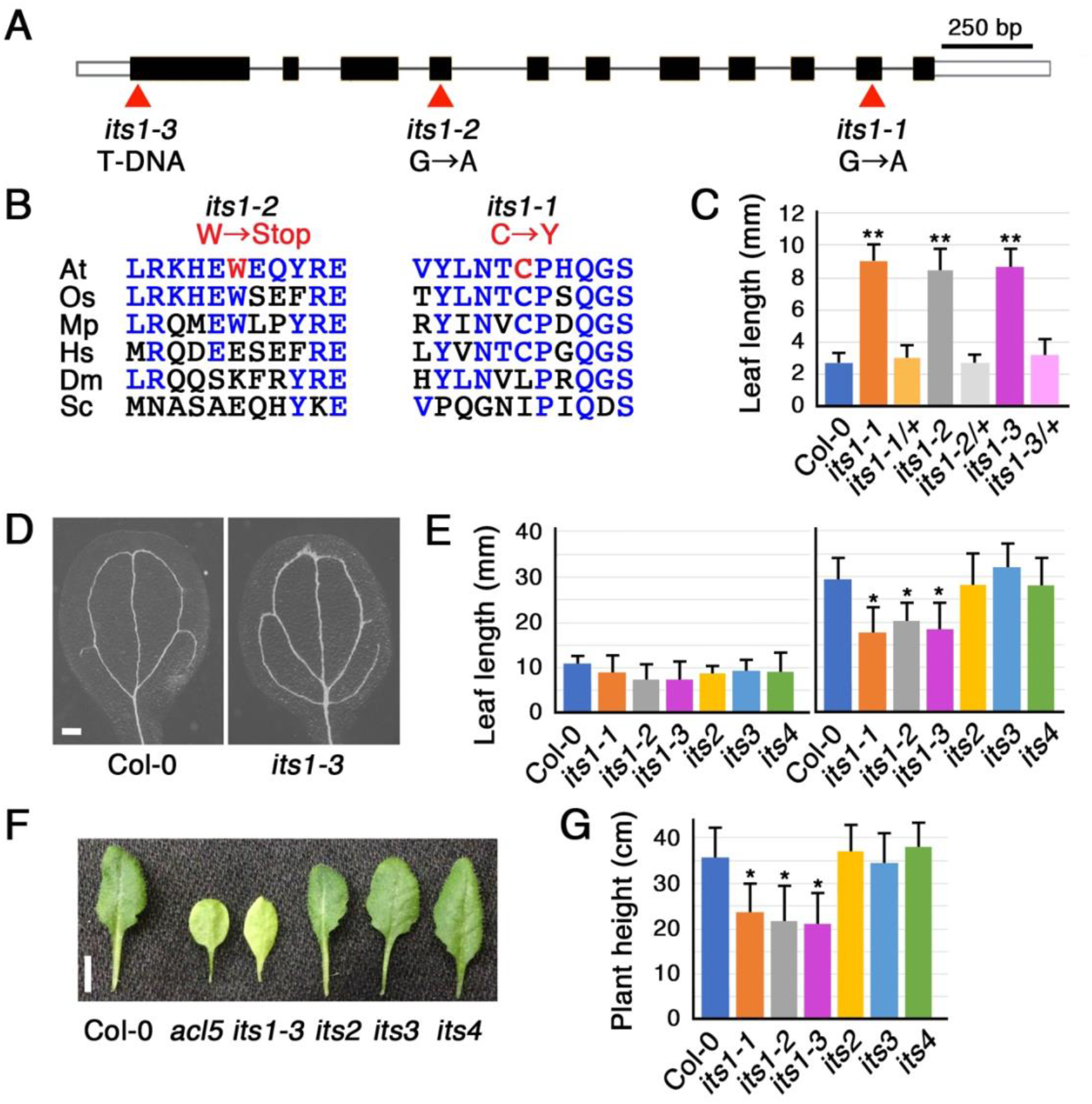
Characterization of an *its1*-responsible gene. A, Gene structure of *ITS1*. Black boxes indicate coding regions and open boxes indicate 5’ leader and 3’ untranslated regions. Mutation sites are indicated by red triangles. B, Alignment of amino acid sequences around the mutation site of *its1-1* and *its1-2* with the corresponding sequences of the homologous proteins of different organisms. Amino acids mutated in *its1-1* and *its1-2* are highlighted in red font. Amino acids identical to those in Arabidopsis are shown in blue font. At, *Arabidopsis thaliana* (NP_197431); Os, *Oryza sativa* (XP_015637029); Mp, *Marchantia polymorpha* (PTQ38793); Hs, *Homo sapiens* (XP_047279415); Dm, *Drosophila melanogaster* (AAM49835); Sc, *Saccharomyces cerevisiae* (NP_011799). C, Comparison of the length of the first leaf between Col-0 and mutant seedlings grown for 7 days with thermospermine. /+ indicates a heterozygote with the wild-type allele. Values are means ± SE (n = 10). Asterisks indicate significant difference from Col-0 (\*\**P* < 0.01, Two-sided Student’s *t* test). D, Vein phenotype of Col-0 and *its1-3* cotyledons. Seedlings were grown for 4 days in agar plates and cleared with chloral hydrate. Scale bar, 200 μm. E, Comparison of the length of the first leaf between Col-0 and mutant seedlings grown for 7 days (left panel) and that of the 5^th^ or 6^th^ leaf between Col-0 and mutant plants grown for 30 days (right panel) in pots. Values are means ± SE (n = 10). Asterisks indicate significant difference from Col-0 (\**P* < 0.05, Two-sided Student’s *t* test). F, Phenotype of the 5^th^ or 6^th^ leaf of Col-0 and mutant plants. Scale bar, 1 cm. G, Comparison of plant height between Col-0 and mutant plants grown for 30 days in pots. Values are means ± SE (n = 10). Asterisks indicate significant difference from Col-0 (\**P* < 0.05, Two-sided Student’s *t* test).

### *its2* represents a weak allele of *CBF5/NAP57*

The *its2* mutant was found to contain a base substitution in *CBF5/NAP57* encoding an rRNA pseudouridine synthase (Figure 3A). In a previous study, two T-DNA insertion alleles *cbf5-1* and *cbf5-2* were shown to be lethal (Lermontova et al., 2007). Because the *its2* mutant shows a recessive inheritance, it was crossed with a heterozygote of another T-DNA insertion allele, which we call here as *cbf5-3* (Figure 3A). The homozygote for this allele was also confirmed to be lethal, previously (Kannan et al., 2008). The resulting F1 seedlings showed segregation of 1:1 in terms of the sensitivity to thermospermine and the insensitive seedlings were confirmed to contain the T-DNA of *cbf5-3* (Figure 3B). Thus, we concluded that *its2* is an allele of *CBF5/NAP57* and name *cbf5-4*. The Gly-to-Arg substitution in *cbf5-4* occurs in the Gly residue highly conserved among plants and animals (Figure 3C). However, in contrast to null mutants, *cbf5-4* plants show normal morphology under standard growth conditions, suggesting that it is a weak loss-of-function mutant.

**Figure 3.**
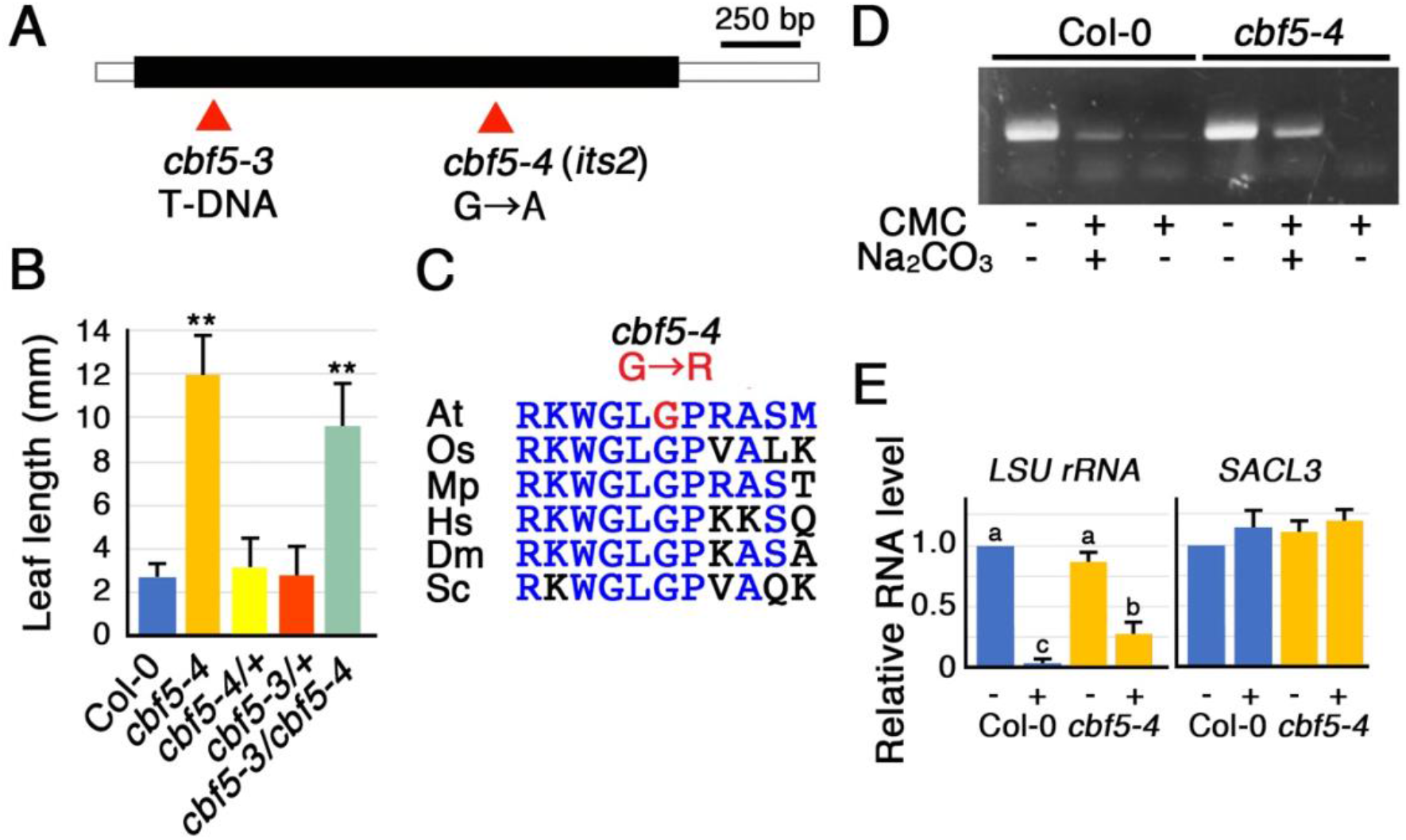
Characterization of an *its2*-responsible gene. A, Gene structure of *CBF5/NAP57*. A black box indicates a coding region and open boxes indicate 5’ leader and 3’ untranslated regions. Mutation sites are indicated by red triangles. B, Comparison of the length of the first leaf between Col-0 and mutant seedlings grown for 7 days with thermospermine. /+ indicates a heterozygote with the wild-type allele. Values are means ± SE (n = 10). Asterisks indicate significant difference from Col-0 (\*\**P* < 0.01, Two-sided Student’s *t* test). C, Alignment of an amino acid sequence around the mutation site of *cbf5-4* with the corresponding sequences of the homologous proteins of different organisms. The amino acid mutated in *cbf5-4* is highlighted in red font. Amino acids identical to those in Arabidopsis are shown in blue font. At, *Arabidopsis thaliana* (NP_191274); Os, *Oryza sativa* (XP_015647556); Mp, *Marchantia polymorpha* (BBN08575); Hs, *Homo sapiens* (NP_001354); Dm, *Drosophila melanogaster* (NP_525120); Sc, *Saccharomyces cerevisiae* (KZV09420). D, Detection of reduced pseudouridylation in the LSU rRNA. Total RNA prepared from Col-0 and *cbf5-4* seedlings was treated with (+) or without (–) CMC, subsequently with (+) or without (–) Na2CO3 as described in the Methods, and reverse transcribed followed by PCR amplification of the LSU rRNA sequence containing Ψ sites and agarose gel electrophoresis. E, Inverse quantification of pseudouridylation in the LSU rRNA. Total RNA was treated with (+) or without (–) CMC, subsequently with Na2CO3, reverse transcribed with rRNA-specific primers for the LSU rRNA or with an oligo(dT) primer for mRNA, and amplified by PCR. Values are means ± SE (n = 6). Different letters indicate significant differences at *P* < 0.05 by ANOVA.

We examined if the level of pseudouridylation in the large subunit (LSU) rRNA is altered or not in *cbf5-4* by using carbodiimide, which specifically reacts with pseudouridine (Ψ) at alkaline pH thereby inhibiting reverse transcription of the RNAs containing Ψ residues (Motorin et al., 2007). Previous studies have reported for Ψ sites in the LSU rRNA in Arabidopsis (Brown et al., 2003; Chen and Wu, 2019; Streit and Schleiff, 2019; Sun et al., 2019). A part of the LSU rRNA containing these Ψ sites (Supplemental Figure S1) was amplified by RT-PCR. The results showed that the level of the cDNA fragment amplified from *cbf5-4* was significantly higher than that from the wild type, suggesting reduced pseudouridylation in *cbf5-4* (Figure 3, D and E). In addition, we examined if the 5’ leader region of the *SACL3* mRNA contains Ψ or not. The cDNA fragment was amplified to the same degree in the wild type as in *cbf5-4*, confirming that this region is not pseudouridylated (Figure 3E).

### *its3* is a semidominant allele of *STIPL1*/*NTR1*

*its3* contains a base substitution in *SPLICEOSOMAL TIMEKEEPER LOCUS1* (*STIPL1*) (Figure 4A), which encodes a homologue of the yeast spliceosome disassembly factor, NTC-Related protein1 (NTR1). However, the region around the amino acid Glu328 replaced with Lys in *its3* may not be conserved among plants, animals, and fungi. The rice gene encodes Lys and another Arabidopsis gene At4g42330 with high homology to *STIPL1* encodes Arg in this position (Figure 4B). F1 progeny and about half of F2 progeny seedlings from a cross of *its3* with the wild type show intermediate sensitivity to thermospermine (Figure 4C), indicating a semi-dominant trait of *its3*. A loss-of-function mutant of *STIPL1, stipl1-1* has been identified as a mutant that induces a long circadian period phenotype under constant conditions and less efficient splicing of circadian-associated transcripts may contribute to the mutant phenotype (Jones et al., 2012). Other studies reported for the involvement of STIPL1 as NTR1 in a transcription elongation checkpoint at alternative exon (Dolata et al., 2015) and in the promotion of miRNA biogenesis (Wang et al., 2019). A T-DNA null allele *ntr1-1*, initially called *stipl1-2*, shows pleiotropic phenotypes including low seed dormancy, delayed flowering, short leaves, and enhanced lethality at high temperatures (Dolata et al., 2015; Wang et al., 2019) but we detected no altered sensitivity to thermospermine in *stipl1-2* (Figure 4C). To confirm whether *its3* is indeed an allele of *STIPL1/NTR1*, we crossed *its3* with *stipl1-2* and found that the F1 seedlings has the same insensitivity to thermospermine as does homozygous *its3* (Figure 4C). We thus concluded that *its3* is a semidominant allele of *STIPL1/NTR1*. Here, considering another T-DNA allele *ntr1-2* (Dolata et al., 2015), we call *its3* as *stipl1-4*. In contrast to temperature-sensitive *stipl1-2, stipl1-4* shows the seedling growth at 30°C comparable to the wild type (Figure 4D). However, as a common phenotype to *stipl1-2, stipl1-4* was shown to have significantly increased and decreased levels of expression of circadian clock genes, *LHY* and *TOC1*, respectively, compared with the wild type at 6 h after 10-day-growth under 16/8-h light/dark cycle (Figure 4E).

**Figure 4.**
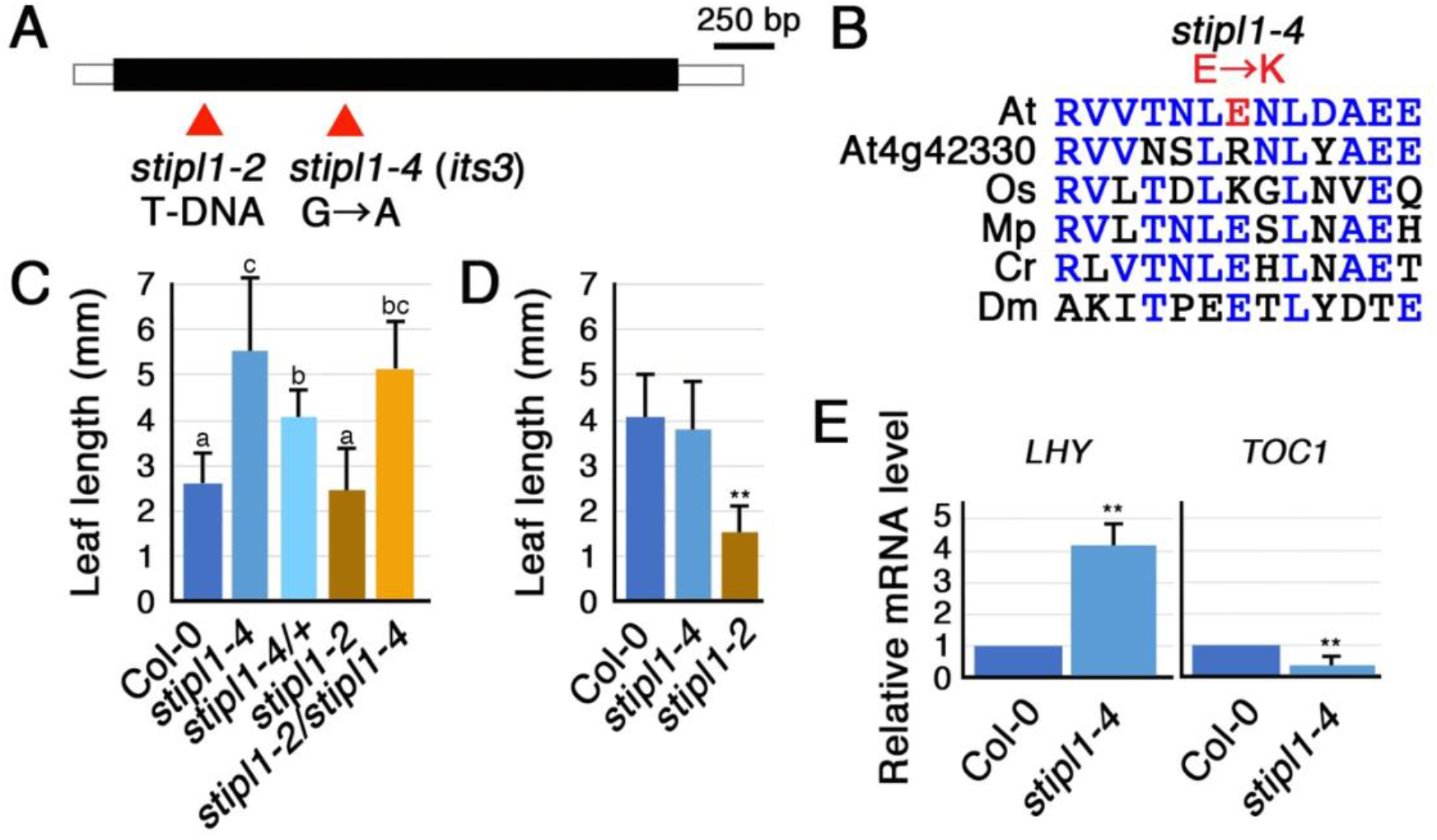
Characterization of an *its3*-responsible gene. A, Gene structure of *STIPL1/NTR1*. A black box indicates a coding region and open boxes indicate 5’ leader and 3’ untranslated regions. Mutation sites are indicated by red triangles. B, Alignment of an amino acid sequence around the mutation site of *stipl1-4* with the corresponding sequences of the homologous proteins of different organisms. The amino acid mutated in *stipl1-4* is highlighted in red font. Amino acids identical to those in Arabidopsis STIPL1/NTR1 are shown in blue font. At, *Arabidopsis thaliana* (Q9SHG6); Os, *Oryza sativa* (BAF30296); Mp, *Marchantia polymorpha* (PTQ30660); Cr, *Chlamydomonas reinhardtii* (PNW80634); Dm, *Drosophila melanogaster* (NP_001285636). C, Comparison of the length of the first leaf between Col-0 and mutant seedlings grown for 7 days with thermospermine. /+ indicates a heterozygote with the wild-type allele. Values are means ± SE (n = 10). Different letters indicate significant differences at *P* < 0.05 by ANOVA. D, Comparison of the length of the first leaf between Col-0 and mutant seedlings grown for 2 days at 22°C and further for 2 days at 30°C. Values are means ± SE (n = 10). Asterisks indicate significant difference from Col-0 (\*\**P* < 0.01, Two-sided Student’s *t* test). E, Effect of *stipl1-4* on the expression of clock genes, *LHY* and *TOC1*. RNA was prepared from Col-0 and *stipl1-4* seedlings at 6 h after 10-day-growth under 16/8-h light/dark cycle and subjected to qRT-PCR. mRNA levels were normalized to the *ACT8* mRNA level and set to 1 in Col-0. Values are means ± SE (n = 6). Asterisks indicate significant difference from Col-0 (\*\**P* < 0.01, Two-sided Student’s *t* test).

### *its4* is a loss-of-function allele of *PHIP1*

Genome mapping and sequence analysis revealed that *its4* is a recessive mutant and has a base substitution in *PHRAGMOPLASTIN INTERACTING PROTEIN1* (*PHIP1*) encoding a plant-specific protein with two RNA recognition motifs (Figure 5A). This mutation introduces a premature stop codon in place of Trp271 within the region conserved among many RNA-binding proteins (Figure 5B). A previous study has identified PHIP1 as a binding partner of phragmoplastin, which is implicated in cell plate formation, and suggests a role of PHIP1 in the polarized mRNA transport to the vicinity of the cell plate (Ma et al., 2008). However, no mutants of *PHIP1* have been characterized. We have tested the sensitivity to thermospermine in a T-DNA insertion mutant of *PHIP1*, which we name *phip1-1*. The *phip1-1* seedling shows a slightly higher insensitivity to thermospermine in the leaf growth (Figure 5C). We then crossed *its4* with *phip1-1* and found that the insensitivity in F1 seedlings is almost like that in homozygous *its4* (Figure 5C). Because the T-DNA in *phip1-1* is inserted near the end of the protein coding sequence, *phip1-1* may be a weak allele. Based on these results, we concluded that *PHIP1* loss-of-function confers thermospermine-insensitive phenotype in *its4*, which we call here as *phip1-2*. These two mutants exhibit no morphological abnormalities under standard growth conditions.

**Figure 5.**
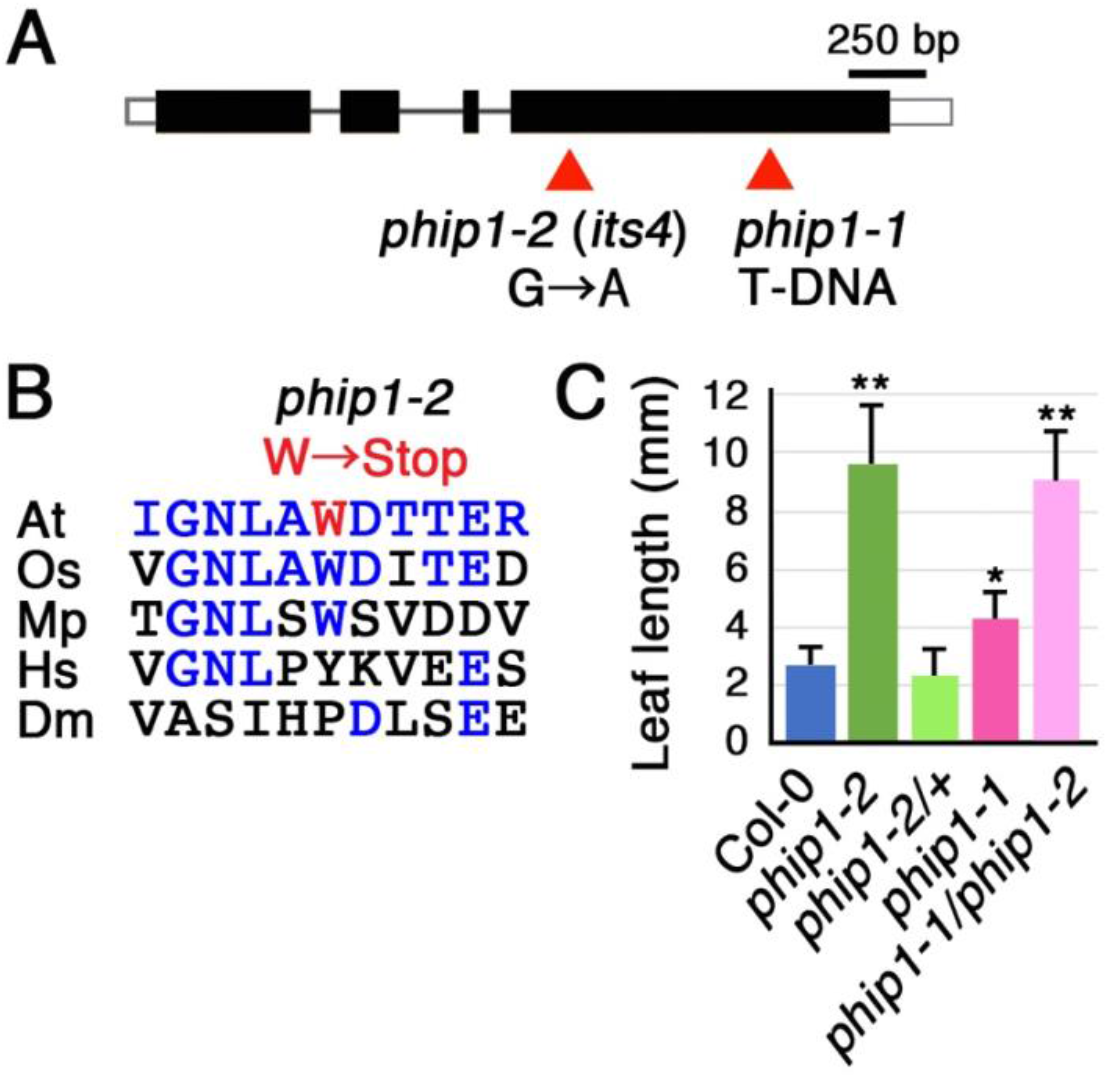
Characterization of an *its4*-responsible gene. A, Gene structure of *PHIP1*. Black boxes indicate coding regions and open boxes indicate 5’ leader and 3’ untranslated regions. Mutation sites are indicated by red triangles. B, Alignment of an amino acid sequence around the mutation site of *phip1-2* with the corresponding sequences of the homologous proteins of plants and those of animals that have only limited homology in RNA-binding domains. The amino acid mutated in *phip1-2* is highlighted in red font. Amino acids identical to those in Arabidopsis are shown in blue font. At, *Arabidopsis thaliana* (OAP06227); Os, *Oryza sativa* (EEE62099); Mp, *Marchantia polymorpha* (PTQ30666); Hs, *Homo sapiens* (NP_055829); Dm, *Drosophila melanogaster* (NP_525123). C, Comparison of the length of the first leaf between Col-0 and mutant seedlings grown for 7 days with thermospermine. /+ indicates a heterozygote with the wild-type allele. Values are means ± SE (n = 10). Asterisks indicate significant difference from Col-0 (\**P* < 0.05; \*\**P* < 0.01, Two-sided Student’s *t* test).

### *its* mutations affect translation of *SAC51* and *SACL3*

Our previous work has revealed that *sac51-1 sacl3-1*double knockout mutants are highly insensitive to thermospermine (Cai et al., 2016). This raises a possibility that expressions of *SAC51* and *SACL3* are affected in *its* mutants. However, the result of qRT-PCR experiments revealed that these mRNA levels are not reduced in *its* mutant seedlings (Figure 6A). *its1-1* seedlings rather have slightly but significantly increased levels of *SACL2* and *SACL3* mRNAs. We then examined the effect of *its* mutations on the mRNA translation of *SAC51* by using transgenic lines carrying a GUS fusion with the *SAC51* promoter and the 5’ leader fragment containing uORFs. Our results revealed that the GUS activities derived from the *SAC51* 5’leader-GUS fusion transcript are severely reduced in *its1-1, cbf5-4* (*its2*), *stipl1-4* (*its3*), and *phip1-2* (*its4*) seedlings (Figure 6B).

**Figure 6.**
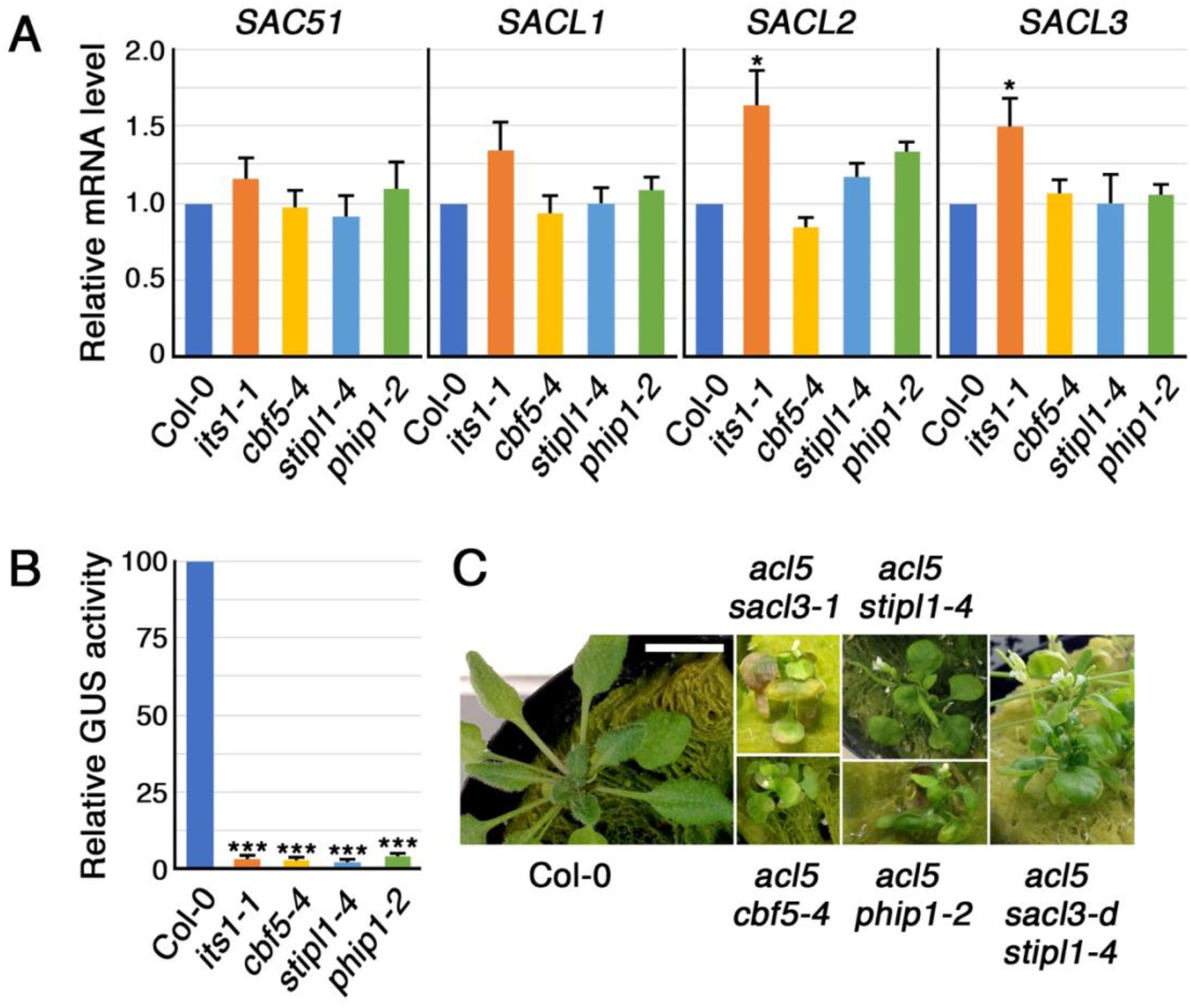
Relationship between *its* mutations and the *SAC51* gene family. A, Effect of *its1-1, cbf5-4, stipl1-4*, and *phip1-2* on the expression of the *SAC51* gene family. RNA was prepared from 10-day-old seedlings of Col-0 and each mutant and subjected to qRT-PCR. mRNA levels were normalized to the *ACT8* mRNA level and set to 1 in Col-0. Values are means ± SE (n = 6). Asterisks indicate significant difference from Col-0 (\**P* < 0.05, Two-sided Student’s *t* test). B, Effect of *its1-1, cbf5-4, stipl1-4*, and *phip1-2* on the GUS activity generated from the *SAC51* promoter-*SAC51* 5’ leader-GUS fusion construct. Protein extracts were prepared from 10-day-old seedlings of each genotype carrying the GUS construct. Values are means ± SE (n = 10). Asterisks indicate significant difference from Col-0 (\*\*\**P* < 0.001, Two-sided Student’s *t* test). C, Phenotype of a 25-day-old plant of Col-0 and 35-to 40-day-plants of each genotype. Scale bar, 1 cm.

As described before, a dominant overexpression allele of *SACL3, sacl3-d*, suppresses the dwarf phenotype of *acl5*, which is defective in thermospermine biosynthesis, while a knockout allele, *sacl3-1*, enhances the *acl5* phenotype, resulting in tiny-sized plants. To examine further the effect of *its* mutations on the *SACL3* mRNA translation, we crossed each *its* mutant with *acl5* and found that double mutants of *acl5 cbf5-4, acl5 stipl1-4*, and *acl5 phip1-2* exhibit tiny-sized plant phenotypes like *acl5 sacl3-1*, although the enhancing effect of *stipl1-4* is moderate compared to that of *cbf5-4* and *phip1-2*, and only *acl5 stipl1-4* plants retain fertility (Figure 6C). So far, we have not obtained *acl5 its1* double mutants because chromosomal locations of *ACL5* and *ITS1* are very close to each other. We have also generated *acl5 sacl3-d stipl1-4* and found that the phenotype of *acl5 stipl1-4* is not suppressed by *sacl3-d*, indicating that the suppressive effect of *sacl3-d* on the *acl5* phenotype is cancelled by *stipl1-4* (Figure 6C).

### Relationship between *its* mutations

To get further information on the function of each gene responsible for *its* mutations, we referred to the co-expression database ATTED II (Obayashi et al., 2007). The list of the top 50 co-expressed genes of *ITS1, CBF5/NAP57, STIPL1*/*NTR1*, and *PHIP1* is shown in Supplemental Table S1. The gene ontology (GO) analysis of these genes confirmed that *ITS1, CBF5/NAP57*, and *PHIP1* are closely associated with the genes involved in rRNA processing and ribosome biogenesis while *STIPL1/NTR1* is linked with those involved in mRNA splicing (Supplemental Table S2). In the list of the genes co-expressed with *ITS1, CBF5/NAP57* is ranked as the 17^th^ and, vice versa, *ITS1* is the 32^th^. *CBF5/NAP57* is also ranked as the 43^rd^ in the list of *PHIP1* co-expressed genes.

We next examined expressions of the four genes in *its* mutants. While the *ITS1* mRNA level is increased in *its1-1* and *cbf5-4*, the *CBF5/NAP57* mRNA level is increased only in *cbf5-4* (Figure 7A). The *STIPL1/NTR1* mRNA level is decreased in *stipl1-4* and *phip1-2*. The *PHIP1* mRNA level is also downregulated in *phip1-2* but upregulated in *cbf5-4*. Expressions of *RPL10A, RACK1A, RPL4A*, and *JMJ22*, identified as a gene responsible for *acl5* suppressors, *sac52-d, sac53-d, sac56-d*, and *sac59*, respectively, were also examined in *its* mutants. *JMJ22* encodes a D6-class Jumonji C (JMJD6) protein implicated in RNA processing (Matsuo et al., 2022). We note that *JMJ22* is ranked the 5^th^ and the 47^th^ in the list of co-expressed genes of *ITS1* and *CBF5/NAP57*, respectively (Supplemental Table S1). The results revealed that only *cbf5-4* has a slight but significant increase in mRNA levels of *RPL10A/SAC52* and *JMJ22/SAC59* (Figure 7B).

**Figure 7.**
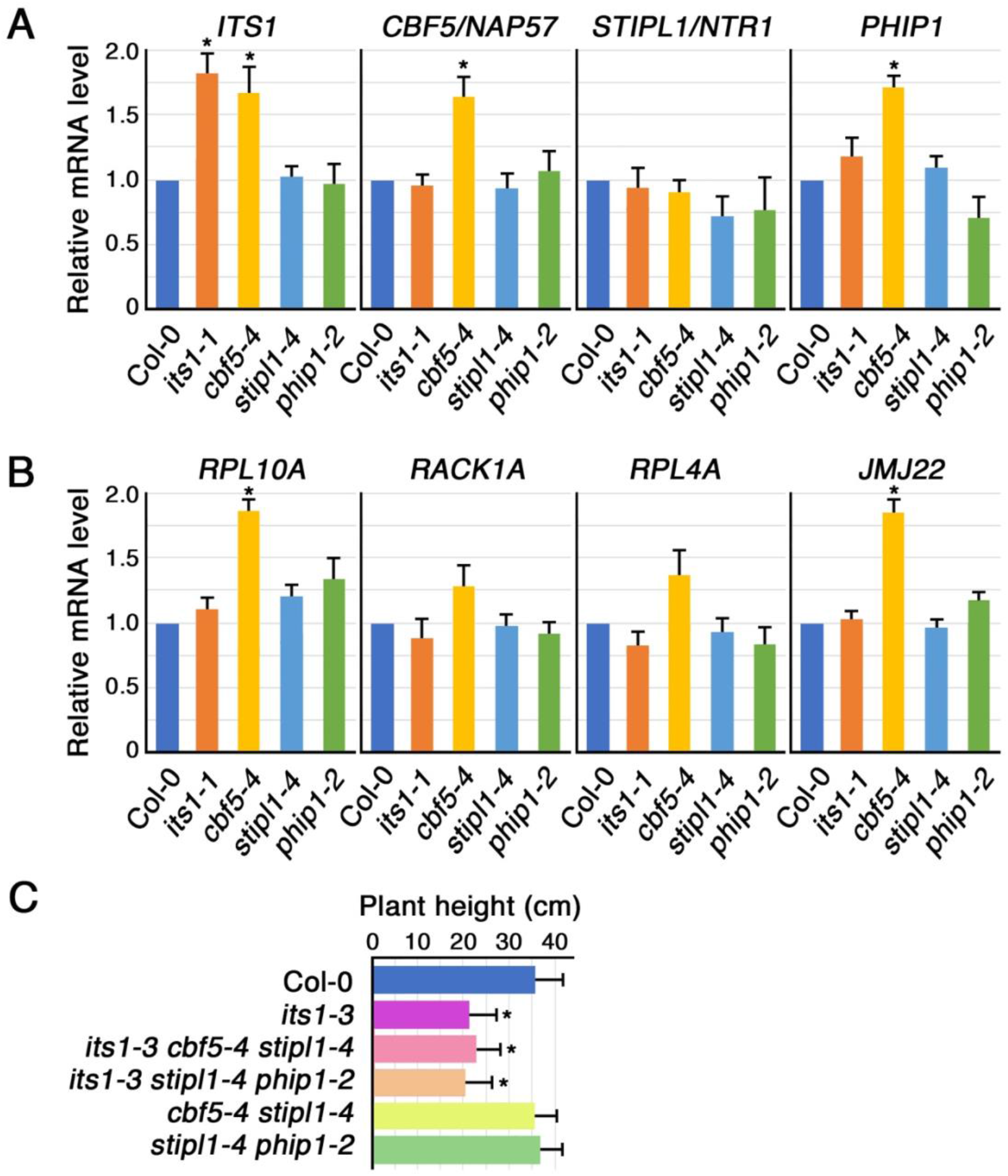
Effect of *its* mutations on their own genes and other *SAC* genes, and effect of *its* multiple mutations on the growth. A and B, Effect of *its1-1, cbf5-4, stipl1-4*, and *phip1-2* on the expression of *ITS1, CBF5/NAP57, STIPL1/NTR1*, and *PHIP1* (A) and that of *RPL10A/SAC52, RACK1A/SAC53, RPL4A/SAC56*, and *JMJ22/SAC59* (B). RNA was prepared from 10-day-old seedlings of Col-0 and each mutant and subjected to qRT-PCR. mRNA levels were normalized to the *ACT8* mRNA level and set to 1 in Col-0. Values are means ± SE (n = 6). Asterisks indicate significant difference from Col-0 (\**P* < 0.05, Two-sided Student’s *t* test). C, Comparison of plant height between Col-0 and mutant plants grown for 30 days in pots. Values are means ± SE (n = 10). Asterisks indicate significant difference from Col-0 (\**P* < 0.05, Two-sided Student’s *t* test).

We generated multiple *its* mutants by crosses. Double mutants of *cbf5-4 stipl1-4* and *stipl1-4 phip1-2* are morphologically normal. Triple mutants of *its1-3 cbf5-4 stipl1-4* and *its1-3 stipl1-4 phip1-2* have no additional phenotype to small leaves and a moderately reduced plant height, which are attributed to *its1-3* (Figure 7C). The *cbf5-4 phip1-2* double mutants have not so far been obtained because of their close chromosomal locations.

## Discussion

In this study, we identified four genes whose mutants exhibit thermospermine-insensitive growth of seedlings and found that all these genes encode proteins related to RNA processing or modification. It is off course possible that further screening for mutants identify additional genes involved in other steps of the response to thermospermine. Our results revealed that mRNA levels of *SAC51* and *SACL3* are not, or almost not, affected but the GUS reporter activity derived from the *SAC51* 5’-GUS fusion is severely reduced in these four mutants. Further, their double mutants with *acl5* exhibit in common the tiny-sized plant phenotype mimicking that of *acl5 sacl3-1*. These findings suggest that all four *its* mutants severely repress the mRNA translation of *SAC51* and *SACL3* and this may cause the thermospermine-insensitive phenotype like that of the *sac51-1 sacl3-1* double knockout. The uORF conserved between *SAC51* and *SACL3* is also present in the 5’ leader of *SACL1* and *SACL2*. Effects of *its* mutations on these mRNAs and those containing a conserved uORF of other classes remain to be investigated.

The growth defects observed in knockout alleles of *ITS1* indicate the functional importance of *ITS1* in other aspects than the response to thermospermine although it is not an essential gene for survival. A recent study has revealed that Upa1 and Upa2, yeast orthologues of SPOUT1/C9orf114, are contained in the primordial pre-60S ribosomal subunit and raised a possibility that they methylate a specific pseudouridine in the pre-60S (Ismail et al., 2022). Multiple mutant alleles of *ITS1* will provide a tool for biochemical characterization of *ITS1* in future research. Although *its1 acl5* has not been obtained because of the close chromosomal position of these two mutations, we speculate that the double knockout would display a tiny-sized plant phenotype with reduced mRNA translation of *SACL3*.

*its2* was found to represent a weak allele of an essential gene *CBF5*. In other organisms, many studies have suggested the significance of Ψ in RNA-protein binding (Wu et al., 2016; deLorimier et al., 2017; Levi and Arava, 2021). In yeast, all rRNA Ψ sites are catalyzed by CBF5 and the CBF5-D95A mutant cells show decreased affinity for tRNAs and decreased translational fidelity (Jack et al., 2011). CBF5/NAP57, which is also called dyskerin, is highly conserved evolutionarily and acts as a core component of a ribonucleoprotein (RNP) complex which associates with RNAs that contain the H/ACA motif. Because of the variety of H/ACA RNAs that guide or specify the target of this RNP complex, this protein has been implicated in diverse processes including ribosome biogenesis, pre-mRNA splicing by binding to small Cajal body (sca) RNAs, and telomere maintenance by binding to the telomerase RNA (Garus et al., 2021). A recent study has shown that the Arabidopsis CBF5 participates in the plant telomerase complex via interactions with the telomerase RNA despite the lack of a canonical H/ACA motif in the telomerase RNA (Song et al., 2021). Although we have examined and detected the effect of *cbf5-4* only on the pseudouridylation of the rRNA, it is possible that this mutation affects the translation of *SAC51, SACL3*, and probably other mRNAs through other pathways involving small nucleolar RNAs. We notice that expression levels of the genes including *CBF5* itself, *ITS1, PHIP1, RPL10A*, and *JMJ22*, are generally upregulated in *cbf5-4*, suggesting a feedback control mechanism that compensates for the defect of the CBF5 function.

On the other hand, our results revealed the involvement of a spliceosome disassembly factor STIPL1/NTR1 in mRNA translation of *SAC51* and *SACL3*. The yeast *NTR1* gene is essential for cell viability and has been shown to associate with a post-splicing complex containing the excised intron and the spliceosomal small nuclear (sn) RNAs (Boon et al., 2006). Recently, TFIP11, the human orthologue of NTR1, has been shown to localize to nucleoli and Cajal Bodies and be essential for the 2’-O-methylation of U6 snRNA, a core catalytic component of the spliceosome, in the nucleolus (Duchemin et al., 2021). Assembly of U4/U6.U5 tri-small ribonucleoproteins is impaired and the fidelity of pre-mRNA splicing is affected by the depletion of TFIP11. The Arabidopsis *stipl1* knockout mutant shows circadian clock defects. We found that the expression of clock genes is affected in the semidominant *stipl1-4* mutant. However, the mutant of a paralogous gene *STIPL2* does not show a circadian phenotype while the double mutants have not been obtained (Jones et al., 2012). Functional difference between these two genes will be elucidated by further investigation with the use of *stipl1-4*.

The *PHIP1* gene responsible for the *its4* mutation was initially identified as and named after a phragmoplastin-interacting protein (Ma et al., 2008). However, no further studies have been reported for *PHIP1*. The *phip1-2* allele contains a premature stop codon in the middle of two RNA binding motifs and may represent a null allele. GO term analysis of co-expressed genes with *PHIP1* strongly suggests the involvement of *PHIP1* in ribosome biogenesis and rRNA processing as well as the relation to *CBF5*. Considering the absence in animals and fungi, it is possible that *PHIP1* acts in a plant-specific RNA modification as-yet-unidentified. Although no phenotype is observed in *phip1-2* under standard growth conditions, detailed studies under various growth conditions might uncover the functional importance of *PHIP1* other than in the response to thermospermine.

In conclusion, our study reveals that RNA modifying and/or processing enzymes play a critical role in the response to thermospermine through activating translation of specific mRNAs containing conserved uORFs. According to the nature of polyamines, thermospermine may directly interact with RNA molecules. We speculate that the activities of some modifying enzymes are dependent on thermospermine, although the reason is unknown why this polyamine is utilized only in plants, in particular the vascular tissue of vascular plants and some bacteria. The effect of thermospermine on modification of RNAs such as rRNA and snRNA should be focused in future experiments.

## Materials and methods

### Plant material and growth conditions

The Arabidopsis (*Arabidopsis thaliana*) accession Columbia (Col-0) was used as the wild type. *acl5-1, sac51-d, sac51-1* (SALK_107954), *sacl3-d*, and sa*cl3-1* (SALK_147291) were as described previously (Cai et al., 2016). *its* mutants were isolated from an ethyl methane sulfonate (EMS)-mutagenized population of Col-0 seeds. Briefly, about 5,000 seeds of Col-0 were treated with 0.2% EMS (Sigma) for 16 h and the M2 progeny seeds were collected as 10 pools of about 500 M1 plants each. About 2,000 seeds from each pool were grown on agar plates supplemented with Murashige-Skoog (MS) nutrients (Wako, Tokyo, Japan), 1% sucrose, and 100 μM thermospermine-4HCl (Santacruz) at 24°C under 16 h light-8 h dark conditions. Putative mutants were selected visually based on the seedling growth. T-DNA insertion mutants of *ITS1* (SALK_202658C), *CBF5/NAP57* (SALK_031065), *STIPL1*/*NTR1* (SALK_073187C), and *PHIP1* (SALK_052754) were obtained from the Arabidopsis Biological Resource Center (www.arabidopsis.org). A transgenic line carrying the *SAC51* promoter-5’ leader-GUS gene fusion construct was as described previously (Ishitsuka et al., 2019). Plants were grown in pots containing rock-wool cubes surrounded with vermiculite at 22°C under 16 h light/8 h dark conditions, except for seedling experiments.

### Mapping and genotyping

For initial mapping of *its* alleles, each mutant was crossed to another wild-type accession Landsberg *erecta* (L*er*). Genomic DNA was extracted from the F2 seedlings that were grown with thermospermine and showed thermospermine-insensitive phenotype. PCR-based mapping was performed with polymorphic markers as described previously (Konieczny and Ausubel, 1993; Tanaka et al., 2020). Each mutation site was identified by whole genome sequencing of the mutant plant DNA using the MGI DNBSEQ-G400 system at Bioengineering Lab. (Sagamihara, Japan).

For generating multiple mutant combinations, genotypes of *acl5-1* and *its* mutant alleles were confirmed by the dCAPS method (Neff et al., 1998). Genotypes of T-DNA insertion alleles were confirmed by PCR using respective gene- and T-DNA-specific primers. Primers and restriction enzymes used are listed in Supplemental Table S3.

### Hormone treatment and microscopy

For ectopic induction of vascular cells in cotyledons, 4-day-old seedlings grown on 1/2 MS plates with 1% sucrose were transferred to liquid 1/2 MS media containing 5% glucose, 5 μM 2,4-D, 1.2 μM kinetin, and 50 μM bikinin with or without 100 μM thermospermine and incubated with gentle shaking for 4 days under 16-h light/8-h dark conditions. Seedlings were fixed in a mixture of ethanol and acetic acid (9:1, v/v) for one day, cleared with chloral hydrate as described (Yoshimoto et al., 2016), and observed under a light microscope equipped with Nomarski DIC optics (DM5000B, Leica, Wetzlar, Germany).

### Gene expression analysis

For RNA preparation, seeds were surface-sterilized in bleach solution containing 0.01% (v/v) Triton X-100 for 3 min, washed 3 times with sterile water, and sown on MS agar plates containing 1% sucrose. Seedlings were grown at 24°C under 16 h light-8 h dark conditions. For thermospermine treatment, 10-day-old seedlings were preincubated for 24 h in liquid MS media with 1% sucrose and further incubated for 24 h with 100 μM thermospermine. Total RNA was prepared by the phenol extraction procedure (Hanzawa et al., 1997) and reverse transcribed using PrimeScript reverse transcriptase (Takara, Kyoto, Japan). Quantitative real-time PCR (qRT-PCR) was performed on three biological replicates using the Thermal Cycler Dice Real Time System TP-760 (Takara) with the Kapa SYBR fast universal qPCR kit (Kapa Biosystems). *ACT8* was used as an internal control. Each sample was duplicated in PCR reactions. Gene-specific primers used for expression analysis are listed in Supplemental Table S4.

### Detection of RNA pseudouridylation

Ψ in rRNA was detected by using a soluble carbodiimide (Motorin et al., 2007). 10 μg of total RNA was dissolved in 30 μL of the reaction buffer (50 mM Bicine, pH 8.0, 7 M Urea, 4 mM EDTA) with 70 mg/ml N-cyclohexyl-N’-β-(4-methylmorpholinium) ethylcarbodiimide (CMC) or without CMC as a control and incubated at 37°C for 20 min. The reaction was stopped on ice by addition of 100 μl of 0.3 M sodium acetate at pH 5.2, 700 μl of ethanol and 1 ul of glycogen at 2 mg/ml. After being precipitated at −80°C, RNA pellets were resuspended in 40 μl of 50 mM Na2CO3 or in Tris-EDTA buffer as a control of no alkaline treatment, incubated at 37°C for 3 h. RNA samples were again precipitated with 1 ml of glycogen solution, 100 μl of 0.3 M sodium acetate, and 700 ul of ethanol at −80°C. Each RNA sample was collected by centrifugation, dried, reverse transcribed with an LSU rRNA-specific primer, 25S-R or 25S-NoPU-R, or an oligo(dT) primer, and amplified by PCR with specific primers (Supplemental Table S4). 25S-NoPU-R primer was used to amplify the LSU rRNA region that does not contain Ψ sites (Supplemental Figure S1) as an internal control for qRT-PCR.

### GUS assay

Fluorometric assay of the GUS activity was performed as described previously (Jefferson et al., 1987). The fluorescence was measured with an RF-5300PC spectrofluorophotometer (Shimadzu, Kyoto, Japan). Total protein content was measured by using the Bradford assay (BioRad).

## Acknowledgements

This work was supported in part by the Japan Society for the Promotion of Science (JSPS) Grants-in-Aid for Scientific Research (Nos. 19K06724) to T.T.

